# Suction feeding performance and prey escape response interact to determine feeding success in larval fish

**DOI:** 10.1101/603316

**Authors:** Noam Sommerfeld, Roi Holzman

## Abstract

The survival of larval marine fishes during early development is strongly dependent on their ability to capture prey. Most larval fish capture prey by expanding their mouth cavity, generating a “suction flow” that draws the prey into their mouth. Larval fish dwell in a hydrodynamic regime of low Reynolds numbers, which has been shown to impede their ability to capture non-evasive prey. However, the marine environment is characterized by an abundance of evasive prey such as Copepods. These organisms can sense the hydrodynamic disturbance created by approaching predators and perform high-acceleration escape maneuvers. Using a 3D high-speed video system, we characterized the interaction between 8-33 day post hatching *Sparus aurata* larvae and prey from a natural zooplankton assemblage that contained evasive prey, and assessed the factors that determine the outcome of these interactions. Larvae showed strong selectivity for large prey that was moving prior to the initialization of the strike. As previously shown in studies with non-evasive prey, larval feeding success increased with increasing Reynolds numbers. However, larval feeding success was also strongly dependent on the prey’s escape response. Feeding success was lower for larger, more evasive prey, indicating that larvae might be challenged in capturing their preferred prey. The kinematics of successful strikes resulted in shorter response time but higher hydrodynamic signature available for the prey. Thus, despite being “noisier”, successful strikes on evasive prey depended on preceding the prey’s escape response. Our results show that larval performance, rather than larval preferences, determines their diet during early development.

## Introduction

The vast majority of marine fishes reproduce by external fertilization, producing small eggs (~1 mm) that drift into the open ocean (Houde 1987; Cowen and Sponaugle 2009; Barneche et al. 2018). Following a brief period of development (usually lasting several days, depending on the ambient temperature) larvae hatch from the egg and begin to feed autonomously (Hunter 1981; Houde 1987; Cowen and Sponaugle 2009). After metamorphosis, the larvae settle into their adult habitat, either pelagic or benthic. This strategy is termed the “bipartite life cycle”, indicating that the planktonic larvae dwell in a habitat that differs from that of the adults (Hunter 1981; Houde 1987; Cowen and Sponaugle 2009). During the pelagic period, larval diets consist of micro- and macro-zooplankton. Similar to many adult fishes, larvae feed by closing the distance to their prey, then lunging towards it while opening their mouth and expanding their buccal cavity. The expansion of the mouth generates a flow of water that carries the prey into the mouth, potentially countering the escape response of the prey (Holzman et al. 2015; China et al. 2017).

During the first few weeks of their lives, larvae of marine fishes experience high mortality rates, eradicating >90% of individuals before they reach metamorphosis. Previous research has identified multiple agents for this mortality, including predation, advection to unsuitable habitats, low food concentrations, and diseases (Hjort 1914; Houde 1987, 2008). However, the hydrodynamic environment in which larvae dwell also impedes their feeding performance, leading to reduced feeding success, low feeding rates, and possibly starvation (China and Holzman 2014; Yavno and Holzman 2017; Koch et al. 2019). In general, the interaction between a solid (e.g. a prey) and the flow around it (e.g. the suction flow of a feeding fish) can be characterized by the dimensionless Reynolds number (Re), depicting the ratio between inertial and viscous forces exerted on the solid particle (Vogel 1994). Larger objects in faster flows are characterized by a hydrodynamic environment of high Re, in which inertial forces dominate and flows can be turbulent. Smaller objects (such as zooplankton) in slower flows (such as the suction flows of larval fish) are characterized by a hydrodynamic environment of low Re, in which viscous forces dominate, and the flows can be laminar and reversible. Successful feeding events of *Sparus aurata* larvae on rotifers have been characterized by higher Re numbers compared to unsuccessful attempts (China and Holzman 2014; China et al. 2017). The increase in Re has been positively correlated with larval length, and mechanistically attributed to successful larvae expanding their buccal cavity faster (resulting in faster suction flows), and opening their mouth to a larger diameter (China and Holzman 2014; China et al. 2017). While much is known about the interaction between larval fish and inert prey (Hernández 2000; Krebs and Turingan 2003; China and Holzman 2014; China et al. 2017), this knowledge offers only a limited insight into the interaction in nature, in which potential prey species usually possess a high ability to sense and respond to approaching predators.

Copepods are often the dominant zooplankton within the pelagic habitat, and an important food source for both adult fish and their larvae. Marine pelagic copepods are highly sensitive to hydrodynamic disturbances, which they perceive via the movement of small sensory setae located on their antennae (Yen et al. 1992; Yen and Strickler 1996; Fields and Yen 1997). The setae bend under shear that may be generated by the movements of organisms (both predators and prey) near the copepod. Strong shear usually trigger an extremely fast escape response, in which a copepod can accelerate at ~300 ms^−2^ and speeds of ~ 0.5 ms^−1^ (Buskey and Hartline 2003; Strickler and Balázsi 2007). Both sensory and motor capacities of copepods improve over ontogeny, leading to a more efficient escape response.

In *Acarcia tonsa* and *Temora longicornis*, adult copepods were ~6 fold more sensitive than the nauplii (Fields and Yen 1997; Titelman 2001). Correspondingly, the capture probability of nauplii into an artificial siphon flow decreased sharply as they matured (Fields and Yen 1997).

It is well established that the ability to capture copepods confers an energetic advantage compared to feeding on other prey types, and that a copepod-based diet increase larval fish survival (Beaugrand et al. 2003; Olivotto et al. 2008; Piccinetti et al. 2014). Stomach content analysis of larval fishes generally reveal a preference for copepods over other prey types, and this preference increases with larval age (Pepin and Penney 1997; Sabatés and Saiz 2000; Fulford et al. 2006; Jackson and Lenz 2016). Such selectivity could result from an ontogenetic shift in larval preferences (i.e. larvae direct more attacks towards copepods as they mature), or from an ontogenetic improvement in larval preference (i.e. larvae experience higher success rates on copepods as they mature), or both. A computational model that calculated the suction forces exerted on escaping prey, predicted that larval ability to counter the prey’s escape force improves dramatically with larval size and age (Yaniv et al. 2014). Accordingly, the diet of clownfish larvae (*Amphiprion ocellaris*) was shown to consist only of copepod nauplii (*Parvocalanus crassirostris*) in the first days post hatching (DPH), and these larvae transitioned to feed on adult copepods only at ~9 DPH (Jackson and Lenz 2016). However, the mechanism behind this pattern is still unclear. Additionally, while the vast majority of marine fishes reproduce by releasing small pelagic eggs and provide no parental care, clownfishes provide parental care for their demersal eggs and their larvae hatch at a relatively large size and developed state (Kavanagh and Alford 2003; Barneche et al. 2018). Therefore, it is unclear how the performance of *Amphiprion* larvae compare with that of the poorly-developed smaller larvae that hatch from pelagic eggs.

Our goal was to characterize the interaction between small pelagic larval fish and prey from a natural zooplankton assemblage that contains evasive prey. Specifically, we sought to (1) determine whether larvae are selective to evasive prey; (2) estimate the variables that affect feeding success on such prey; and (3) characterize the effect of the larvae’s morphology and kinematics on the escape response of the prey. We used 8-33 DPH *Sparus aurata* larvae, as these larvae hatch from small pelagic eggs, representing the common strategy among marine fishes. Larvae and prey were filmed using two synchronized high-speed cameras in a laboratory setup that provided 3D tracking of both prey and predator.

## Methods

### Study organisms

We used gilthead sea-bream larvae (*Sparus aurata* Linnaeus, 1758) as our model for larval feeding (Holzman et al. 2015). *S. aurata* is a pelagic spawner, hatching at ~3.5 mm. Feeding initiates at ~5 DPH at a body length of ~4 mm. Larvae reach the stage of flexion at ~21-24 DPH, at a length of 7-10 mm, depending on conditions. Larvae were provided by the ARDAG commercial nursery (Eilat, Israel). Throughout the experiments larvae were kept at 19°C in aerated seawater at a salinity of 35 ppm. Larvae were obtained prior to daily feeding, hence were food deprived for >12 hrs.

We used a natural assemblage of zooplankton as the prey in all experiments. Prey were obtained by towing a zooplankton net from a boat cruising at low speed, or by a swimmer, depending on the seasonal abundance of zooplankton in the coastal waters of the Gulf of Aqaba, Eilat. Swimmers towed a 1m long, 100 μm zooplankton net with a mouth diameter of ~0.5 m, while the boat towed a longer, 4 m long net. At the end of the tow, the captured zooplankton was sieved through a 500 μm net to remove larger zooplankton, including predatory arrow worms and other elongated organisms, and carefully transferred into a 1 liter holding aerated container until the onset of experiments. A subsample was observed under a stereoscopic microscope to verify that the sample was dominated by copepods; if not, a new sample was obtained. Fresh zooplankton was collected daily for the experiments.

### 3D filming of prey acquisition strikes

We tracked the 3D position of the larvae and their prey during prey acquisition strikes using two synchronized high-speed cameras (Photron Fastcam SA6) operating at 1000 frames per second. Cameras were fitted with Navitar 6000 ultra-zoom lenses, providing 1:3¼ magnification (i.e. a 1 mm long object is projected at 3.25 mm on the sensor) with a depth of field of ~50 mm (Fig 1). Cameras were positioned such that their resolution and magnification were identical. The field of view of each camera was ~ 40 mm × 30 mm (W × H) at a resolution of 1920 × 1440 pixels. The cameras were positioned 45 cm apart, at an angle of 35° relative to one another (Fig 1). The volume on which the two cameras were focused was ~20 milliliters. To minimize reflections and distortions and maximize the depth of field, the aquarium was constructed such that each phase was perpendicular to one camera. Two rectangular 2.2 watt LED lights were positioned behind the aquarium, providing a backlight illumination of the visualized volume. Reconstruction of points in the 3D space was done using the package DLTdv5 in MATLAB (Hedrick 2008). The system was calibrated at the beginning of each recording session using a calibration grid of 60 points, spanning the visualized volume. Accuracy was assessed by measuring four known distances in three different images, and estimated as <1.5%.

**Fig 1:**
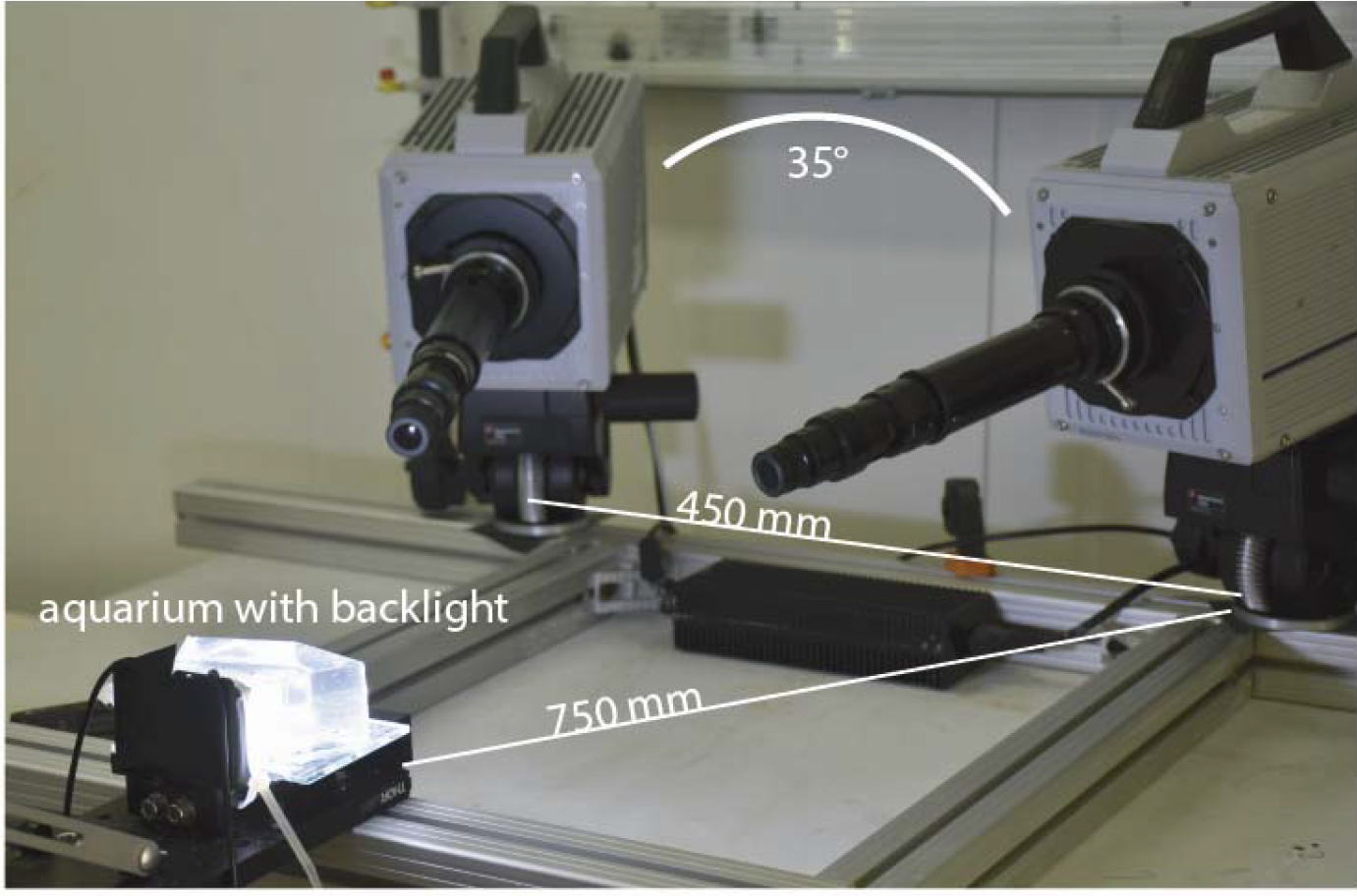
The positions of the larvae and their prey were tracked in 3D using two synchronized high-speed cameras (Photron Fastcam SA6) fitted with Navitar 6000 ultra-zoom lenses. The lenses provided 1:3¼ magnification with a depth of field of ~50 mm. Data were collected at 1000 frames per second.

For each filming session, 5-15 larvae were introduced into the filming chamber and given several minutes to acclimate. A random assortment of prey was then introduced into the chamber using a pipette, and the fish were allowed to feed freely for ~30 min. The system was triggered manually upon the observer’s detection of a predator’s feeding attempt or a prey’s escape response in the visualized volume. Thus, our dataset included clips that featured: (1) predatory strikes in which the prey initiated an escape response; (2) predatory strikes in which the prey did not move; and (3) escape responses executed by the prey before the predator opened their mouth. For each strike, the time of strike initiation (t=0) was defined as the time at which the prey started to escape (cases 1 & 3 above) or as the time when the larvae opened its mouth (case 2). Note that these times were highly correlated (r=0.75) in case 1. Overall, these events involved ~100 larvae ranging in age from 8 to 33 DPH (3-20 mm; Fig 1).

For the two views of each recorded event (from the two cameras that comprise the system) four landmarks were digitized in each frame: the anterior tip of the upper jaw, the anterior tip of the lower jaw, a point on the body (center of the eye), and a point on the prey denoting the edge closest to the fish’s mouth. Four additional landmarks were digitized in one of the frames: (1) a point on the prey denoting the edge furthest from the fish’s mouth, (2) the base of the larva’s caudal fin (3 & 4) the two vertices on the horizontal major axis of an imaginary ellipse encapsulating the prey. Digitized 2D coordinates of the landmarks from the paired cameras were converted to an earthbound 3D coordinate system using DLTdv5. We used the coordinates of the landmarks to calculate the following variables: (1) larval length, calculated as the distance between the center of the mouth to the base of the caudal fin; (2) mouth gape (mm; hereafter “gape”), calculated at each point in time as the distance between the anterior tip of the upper and lower jaw; (3) time to peak gape (TTPG; ms), calculated as the time it took the larva to open its mouth to 95% of maximal gape diameter; (4) gape opening speed (mm/s), calculated as the derivative of gape diameter with time; (5) response distance (mm), the distance of the prey from mouth center at the time of strike initiation; (6) larval swimming speed (mm/s) calculated as the average speed of the larva during the feeding attempt; (7) the time to prey capture (ms); (8) prey cruising speed (mm/s), calculated based on the displacement of the prey 10 frames before the predator’s strike (of prey escape) was initiated; (9) prey escape speed (mm/s), calculated based on the displacement of the prey during its escape, usually <10 frames; and (10) prey length (mm). We used these values to calculate the Reynolds number (equation 1) for feeding and swimming of the larvae. Re was calculated as

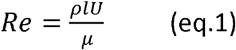

where ρ is the density (1024 kg m^−3^) and μ is the dynamic viscosity (Pa s^−1^) of the fluid. Re_(feeding)_ was calculated using maximal gape diameter as the relevant length (*l*; m) and the peak suction flow speed (*U*; ms^−1^). The latter speed was estimated based on TTPG, maximal gape, and estimated buccal dimensions (Yaniv et al. 2014; China et al. 2017). Re_(swimming)_ was calculated using the larva length as the relevant length (*l*; m), and the larval swimming speed as the flow velocity (*U*; ms^−1^).

### Determinants of feeding success

We used logistic regression to test the effect of the kinematic and morphological traits of both prey and predator on feeding success. The dependent variable was larval feeding success (fail *vs* success; binary variable). The independent variables were selected following China et al (China et al. 2017), who found that the variance in feeding success on non-evasive prey was explained only by the hydrodynamic environment (Re). Because our prey was evasive, we also included variables that can affect the prey’s performance, i.e. whether it initiated an escape response (yes/no; binary variable), prey length, response distance, and prey swimming speed (before the strike was initiated). We used a model averaging approach to identify and weight the variables that affect feeding success. We used the function dredge (Barto’n 2017) to identify the best supported models (with ΔAIC < 2 relative to the best model), followed by calculation of model averaged estimates of the effect size and standard error for each variable (Claeskens and Hjort 2008).

We used a logistic regression model to test which variables determined the prey’s escape response. Copepods are known to execute an escape response when exposed to high strain rates. These can be caused by the body of the approaching predator, in which case the disturbance is expected to increase with the swimming speed of the predator, the radius of the fish’s head, and the distance between the predator and prey. Kiørboe (Kiørboe 2008) estimated this disturbance (γ; s^−1^) as

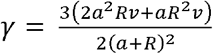

where *R* is the distance between the anterior end of the larvae and the prey (m), *a* is the radius of the larva’s head (m), and ν is the swimming speed of the larvae (ms^−1^). Our model therefore included prey length, larval swimming speed, the radius of the larva’s head, and strike initiation distance as independent variable, and prey escape as a binary dependent variable.

### Selectivity

We characterized the prey available to the larvae before each strike by measuring the length and tracking the motion of all potential prey items located within the larvae’s reactive volume. That volume was defined as a hemisphere with a diameter of 1 larval body length, centered at the larvae’s mouth, and with the plane passing through the center of the hemisphere perpendicular to the larva’s long axis (Fig 2). We used only strikes in which the reactive volume contained more than one prey (N=90). We tested the effect of the length (mm) and motion (binary variable: moving/stationary) on the probability of a larvae striking a prey item using conditional logistic regression. In this analysis, each strike was considered a choice experiment, incorporated into the model as a stratum [i.e. random variable; (Aizaki and Nishimura 2008)].

**Fig 2.**
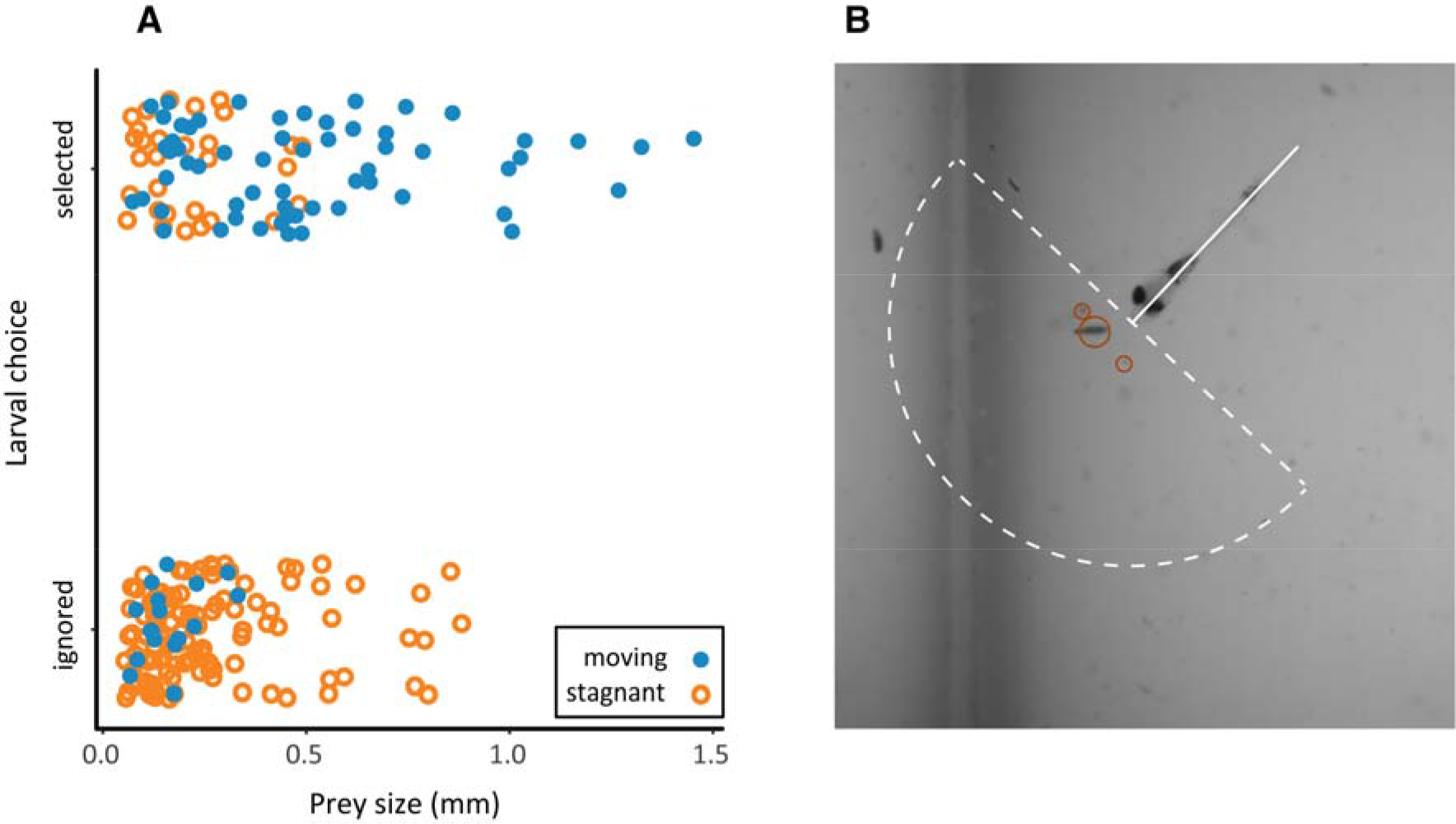
Larvae tended to strike larger, moving prey (Panel A; Table 1). Point colors depict whether the prey was static (orange) or moving (blue) in the 10 frames preceding the strike. Data refer to zooplankton located within an imaginary hemisphere (white dashed line), with a diameter of 1 larval body length, centered at the larvae’s mouth (Panel B).

## Results

### Selectivity

In all strikes in which more than one prey was present at a distance of 1 larval length, attacks were directed towards the larger, moving prey (conditional logistic regression; P<0.001 for size and P<0.007 for movement; whole model R2 = 0.26, P<0.001; N=90 strikes and 223 prey items; Fig 2; Table 1). The logistic regression indicated that a 1 mm increase in the size of the prey would double the probability of attack, and that prey movement would increase attack probability by four-fold (Table 1). Accordingly, the size of attacked prey ranged from 0.06 to 1.5 mm (median 0.36 mm) and that of ignored prey ranged 0.06-0.9 mm (median 0.18). Roughly two-thirds of the attacks were directed at prey that were moving before the strike was initiated (Fig 2).

**Table 1:** Conditional logistic regression model depicting the effect of prey’s length (mm) and motion (binary variable: moving/stationary) on the probability of a fish attempting to capture it. SE- adjusted standard error of the estimate.

|  | Estimate | SE | Z value | P |
| --- | --- | --- | --- | --- |
| Size | 4.07 | 1.50 | 2.71 | 0.007 |
| Motion | 2.17 | 0.43 | 5.05 | 0.0001 |

### Strike success

A model selection procedure identified seven models that best predict strike outcome based on the independent variables (Table 2). The R^2^ for the models ranged from 0.54 to 0.59. A model averaging procedure on the selected models revealed that prey capture significantly increased when prey did not attempt an escape response, when prey length was smaller, and when Re_(feeding)_ was higher (all P<0.05; Table 2; Fig 3). This procedure also provides the relative importance values of each term, calculated as a sum of the Akaike weights over all of the models in which the term appears. The absence of an escape response was the most important predictor of strike success (relative rank =1), followed by Re_(feeding)_ (0.9), Re_(swimming)_ (0.89), prey length (0.8), strike initiation distance (0.56), and prey swimming speed (0.43).

**Table 2:** Conditional average from the seven best logistic regression models depicting the effect of independent variable on feeding success. SE- adjusted standard error of the estimate. Rank is the relative importance values of each term, calculated as a sum of the Akaike weights over all of the models in which the term appears.

|  | Estimate | SE | Z value | P | Rank |
| --- | --- | --- | --- | --- | --- |
| Intercept | 1.31 | 0.86 | 1.52 | 0.127 |  |
| Prey escape (Y/N) | -2.04 | 0.77 | 2.66 | 0.008 | 1 |
| Prey length | -6.66 | 2.97 | 2.23 | 0.025 | 0.8 |
| Re (feeding) | 0.04 | 0.016 | 2.11 | 0.034 | 0.9 |
| Re (swimming) | 0.004 | 0.002 | 1.75 | 0.080 | 0.89 |
| Strike initiation distance | -2.78 | 1.80 | 1.54 | 0.124 | 0.56 |
| Prey speed | -0.095 | 0.080 | 1.18 | 0.238 | 0.43 |

**Fig 3.**
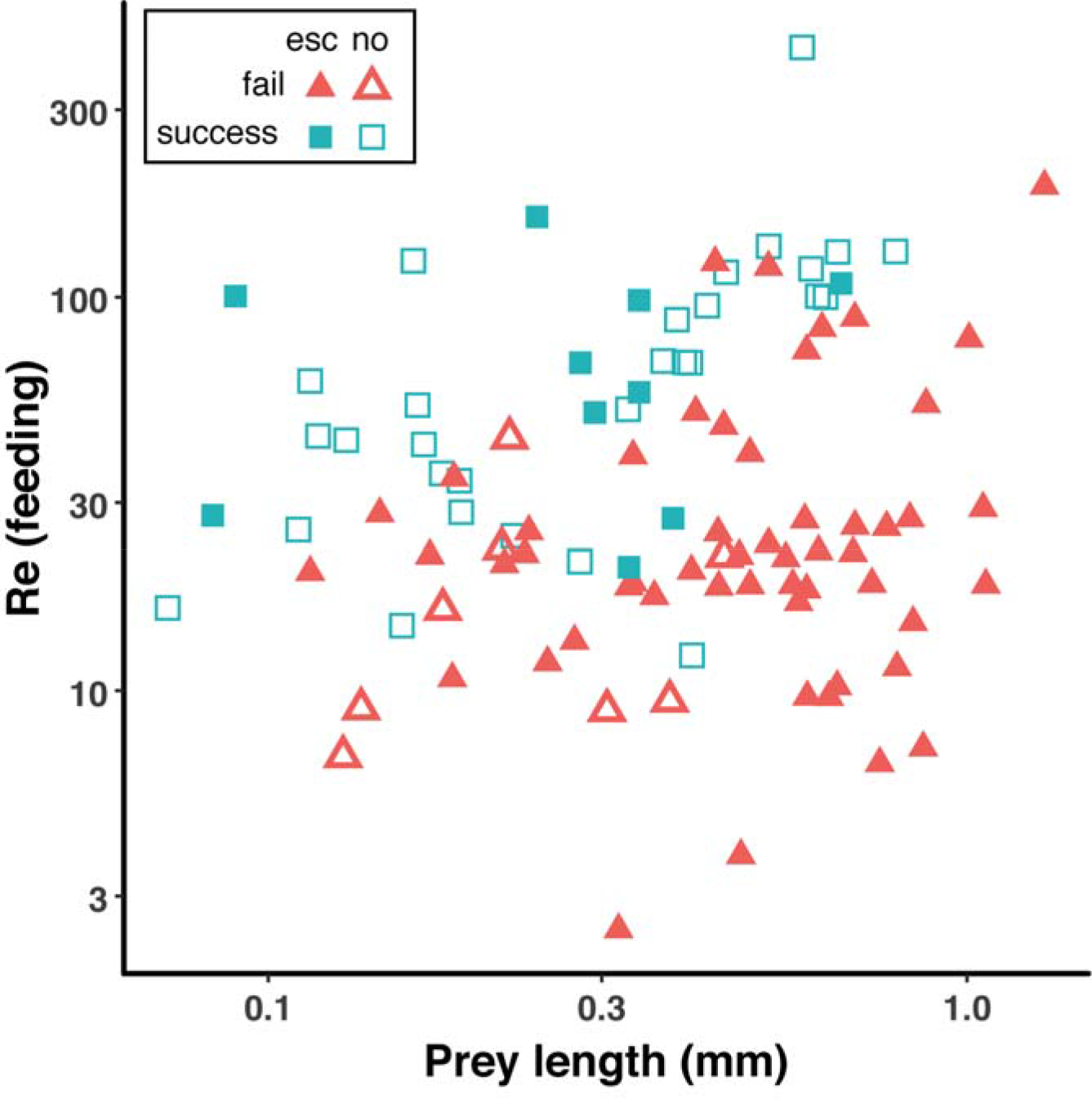
Prey capture success by larval *S. aurata* increased with Re numbers, and decreased for larger prey that elicited an escape response (Table 2). Re_(feeding)_ refers to the Re calculated based on mouth diameter and estimated suction flow speed. Open symbols denote strikes in which the prey did not escape, and full symbols denote strikes in which the prey elicited an escape response. Red triangles indicate larval failure to capture their prey, green squares indicate success. Both X- and Y- axes are plotted on a logarithmic scale.

### Prey escape response

Prey escape response was the most important factor that determined the feeding success (see above). We therefore used a logistic regression model to test which factors affect the probability of the prey producing such a response. The model indicated that increasing larval speed and decreasing strike initiation distance significantly reduced the probability of escape response by the prey (P < 0.006 for both, R2 = 0.42; Table 3; Fig 4), whereas the effect of the other variables was non-significant. Thus, the response time available for the prey (strike initiation distance divided by larval speed) was shorter for prey that did not escape (mean ± SE = 0.01 ± 0.0025 seconds) than for prey that did escape (0.07 ± 0.01 seconds; Fig 4B). Additionally, the mean hydrodynamic disturbance (γ, s^−1^ was lower for prey that did escape than for prey that did not escape (mean ± SE = 12.3 ± 2.0 s^−1^ and 21.2 ± 2.4 s^−1^ for escaping and non-escaping prey, respectively; Fig 4C).

**Table 3:** Logistic regression model depicting the effect of the independent variable on prey escape response. The model’s R^2^ was 0.42. SE- adjusted standard error of the estimate.

|  | Estimate | SE | Z value | P |
| --- | --- | --- | --- | --- |
| Intercept | -0.43 | 0.90 | -0.47 | 0.63 |
| Prey length | 2.50 | 1.73 | 1.44 | 0.14 |
| Larval speed | -0.04 | 0.01 | -4.46 | <0.001 |
| Head radius | 0.32 | 2.78 | 0.11 | 0.91 |
| Strike initiation distance | 3.24 | 1.18 | 2.74 | 0.006 |

**Fig 4.**
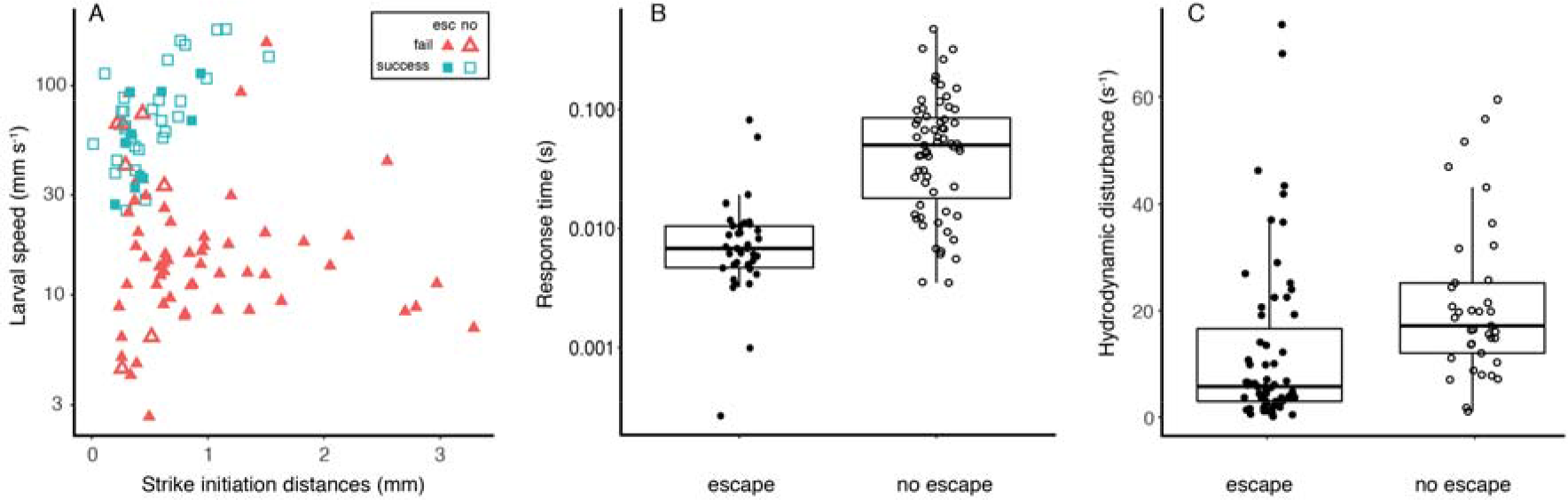
The probability of the prey executing an escape response increased at slower larval swimming speeds and in strikes with longer initiation distances (Panel A; Table 3). Consequently, the response time available for the prey (panel B) was shorter for prey that did not escape (mean ± SE = 0.01 ± 0.0025) than for prey that escaped (mean ± SE = 0.07 ± 0.01). The calculated hydrodynamic disturbance (panel C) was higher for prey that did not escape (mean ± SE = 21.2 ± 2.4 s^−1^) than for prey that produced an escape response (12.3 ± 2.0 s^−1^). Open symbols denote strikes in which the prey did not escape, full symbols denote strikes in which the prey produced an escape response. Red triangles in A indicate larval failure to capture their prey, green squares indicate success. Boxes denote the 1^st^ and 3^rd^ quartiles, black horizontal line is the median, and whiskers denote 1.5 inter-quartile range. Note the logarithmic scale for the Y-axis in A and B.

## Discussion

In this study, we characterized the interaction between larval fish and prey present in a natural zooplankton assembly that was dominated by evasive prey. Larvae showed a strong selectivity for large prey that were moving prior to the initialization of the larva’s strike (Fig 2). As previously shown in studies with non-evasive prey, we found that larval feeding success increased with increasing Reynolds numbers (Fig 3). However, larval feeding success was also strongly dependent on the prey’s escape response (Table 2). Feeding success was lower for larger, more evasive prey (Fig 3), indicating that larvae might be challenged in capturing their preferred prey. The kinematics of strikes on escaping prey were characterized by slower larval swimming speed and greater strike initiation distance compared to strikes on non-escaping prey (Fig 4; Table 3). These kinematics resulted in shorter response time and higher hydrodynamic disturbance for prey that did not escape (Fig 4).

Previous studies of feeding success by larval fish have focused on their interactions with non-evasive prey (Hernández 2000; Krebs and Turingan 2003; China and Holzman 2014; China et al. 2017) although several studies have reported on interactions between copepods and clownfish larvae (Jackson and Lenz 2016; Robinson et al. 2019; Tuttle et al. 2019). In small marine larvae that hatch from pelagic eggs, the hydrodynamic environment (denoted by Re) is the dominant factor that determines larval kinematics and prey capture performance. However, clownfish larvae hatch at a well-developed state following parental care of the eggs, and likely live in a realm of higher Re. It is therefore unclear how their interactions with evasive prey represent the general case across larvae of marine fish, and a direct comparison of the predatory strategies of representative larvae from these two life-history strategies is warranted.

In general, the dynamics of the predator-prey interactions can change due to the prey’s escape response. A CFD model of larval suction flows showed that smaller (younger) larvae would be able to capture only weakly evasive prey that are attacked from a short distance [relative to their mouth diameter (Yaniv et al. 2014)]. The predictions of that CFD model for inert prey were the opposite: that the distance in which such prey could be captured decreased through ontogeny (Yaniv et al. 2014). Observations on *Amphiprion ocellaris* larvae feeding on the calanoid copepod *Bestiolina similis* (Jackson and Lenz 2016; Tuttle et al. 2019) revealed that feeding success on evasive prey increased throughout ontogeny, and that older larvae were able to capture more evasive prey and from a greater distance compared to younger larvae. The results of the present study (Fig 4A) further corroborate the CFD prediction (Yaniv et al. 2014), and demonstrate that for interactions with evasive prey, Reynolds number is not the only parameter that determines strike success (Table 2). Specifically, the ability of the prey to execute an escape response at the right time is critically important in determining the outcome of this predator-prey interaction (Table 2). In predator-prey interactions between adult zebrafish and their prey (larval zebrafish), prey that did not initiated an escape response were always captured, whereas escape responses that were timed correctly resulted in prey escape (Stewart et al. 2013). In the present study, prey that did not escape were not always captured, likely due to the larva’s inability to produce sufficiently strong suction flows. Thus, in larval fish, prey capture is determined on the one hand by the ability of the larva to execute a high-Re strike; and on the other hand by the ability of the prey to execute a timely escape response (Fig 3, 4).

Copepods are well-known for their ability to execute high-acceleration escape responses when sensing a hydrodynamic disturbances (Yen et al. 1992; Fields and Yen 1997; Buskey et al. 2002; Tuttle et al. 2019). Viitasalo et al (Viitasalo et al. 1998) assessed the factors that determine the success of predatory strikes by adult three-spine sticklebacks on two species of copepods. They concluded that feeding success was limited to cases in which the fish was able to approach the copepod slowly. In those cases, the hydrodynamic signal available to the copepod was weaker, resulting in a late escape response and a shorter reaction distance to the approaching predator (Viitasalo et al. 1998). Similar results were observed for *Amphiprion ocellaris* larvae feeding on the calanoid copepod *Bestiolina similis* (Tuttle et al. 2019). In the present study we found an opposite trend: feeding success was limited to cases in which the larvae swam quickly towards the prey. Similar to sticklebacks, prey capture success was associated with short reaction distances. In *S. aurata* larvae, the strike kinematics on prey that eventually executed an escape response resulted in a longer response time and lower hydrodynamic disturbance (Fig 4). We suggest that this trend might reflect the larvae’s inability to correctly time their strike. Striking from too far would allow evasive prey long enough time (~70 ms; Fig 4B) to respond to the hydrodynamic disturbance produced by the predator. Conversely, a stealth approach followed by a fast lounge might provide the prey with little time (<10 ms) to respond to the predator (Tuttle et al. 2019). Thus, despite being “noisier”, successful strikes on evasive prey might depend on striking fast to precede the prey’s escape response. Alternatively, it could also be that the predators are able to distinguish weakly evasive prey and change their kinematics accordingly.

In general, adult fishes show a strong selectivity for large prey (O’Brien et al. 1976; Gardner 1981; Li et al. 1985; Holzman and Genin 2003, 2005). Werner and Hall (Werner and Hall 1974) suggested that such size selection is related to the optimal allocation of time spent searching and handling prey. In contrast, larval fish are considered selective for smaller, less evasive prey, at least in the first days after exogeneous feeding initiates (Pepin and Penney 1997; Sabatés and Saiz 2000; Fulford et al. 2006; Jackson and Lenz 2016). It is nevertheless unclear why selectivity for small prey might be optimal for larvae. However, studies on larval fish selectivity have largely been based on assessing the depletion of prey within an experimental container, or on a comparison between the prey found within the guts of larvae and that in the environment. Either way, such studies integrate two processes within the predator-prey interaction: the first being the recognition and approach to the prey; and the second being the strike itself. In the first stage, selectivity can develop following a bias towards a preferred prey, or due to a difference in prey detectability, with both resulting in a higher attempt rate on a certain prey type (Werner and Hall 1974; Li et al. 1985; Buskey et al. 1993; Holzman and Genin 2005). In the second stage, selectivity can develop following a bias in the ability of the predator to capture certain prey types that that better escape or defend themselves. Our experimental system provides novel insights into the role of each stage in determining larval selectivity. Larvae show a strong preference for directing predatory strikes towards larger, moving prey (Fig 2). However, this preference would not be reflected in their diet, because such prey is more likely to successfully escape (Fig 2, 3). Thus, the apparent selectivity for smaller prey by younger fish larvae could be the result of a prey-size-dependent capture success rather than an active preference for smaller prey.

